# Geographic variation in reproductive assurance of *Clarkia pulchella*

**DOI:** 10.1101/372375

**Authors:** Megan Bontrager, Christopher D. Muir, Amy L. Angert

## Abstract

Climate can affect plant populations through direct effects on physiology and fitness, and through indirect effects on their relationships with pollinating mutualists. We therefore expect that geographic variation in climate might lead to variation in plant mating systems. Biogeographic processes, such as range expansion, can also contribute to geographic patterns in mating system traits. We manipulated pollinator access to plants in eight sites spanning the geographic range of *Clarkia pulchella* to investigate geographic and climatic drivers of fruit production and seed set in the absence of pollinators (reproductive assurance). We examined how reproductive assurance and fruit production varied with the position of sites within the range of the species and with temperature and precipitation. We found that reproductive assurance in *C. pulchella* was greatest in populations in the northern part of the species’ range, and was not well-explained by any of the climate variables that we considered. In the absence of pollinators, some populations of *C. pulchella* have the capacity to increase fruit production, perhaps through resource reallocation, but this response is climate-dependent. Pollinators are important for reproduction in this species, and recruitment is sensitive to seed input. The degree of autonomous self-pollination that is possible in populations of this mixed-mating species may be shaped by historic biogeographic processes or variation in plant and pollinator community composition rather than variation in climate.

## Introduction

The reproductive success of primarily outcrossing plant taxa is often highly dependent on the actions of their mutualist pollinators (Burd, 1994; Ashman et al., 2004). Populations may vary in the degree to which they require pollinators as an adaptive response to mate or pollen limitation, or as a result of historic biogeographic processes (Busch, 2005). Understanding how populations vary in their reliance on pollinator service is important in an era of global change, as changes in phenological cues might lead to mismatch between plants and pollinators (Kudo and Ida, 2013), pollinator populations may decline if they are maladapted to changing conditions (Williams et al., 2007), and climate stress can affect plants’ ability to gather and allocate resources for pollinator attraction and reward (Mu et al., 2015).

In the face of sustained mate or resource limitation, reliance on outcross pollen can limit seed production, and selection might favor individuals with floral traits that facilitate reproductive assurance via self-pollination (Bodbyl Roels and Kelly, 2011). Reproductive assurance is the ability to self-pollinate in the absence of pollinators in order to offset deficits in pollen delivery (Jain, 1976). Limited resources, including limited water availability, can increase the cost of producing and maintaining attractive floral displays (Galen et al., 1999). This could lead to selection for individuals that can achieve high reproductive success without incurring the costs of showy displays. Similarly, short flowering seasons may increase the risks of waiting for pollinator service. Mate limitation can occur when populations are small or sparse, or when pollinators are low in abundance or prefer to visit co-occurring species (Knight et al., 2005). These patterns of mate and resource limitation can co-vary with climatic conditions. Therefore, patterns in mating system traits may be correlated with the climatic gradients that underlie a species’ geographic distribution (Lennartsson, 2002; Moeller and Geber, 2005) and may covary with geographic range position. These climatic gradients may simultaneously affect other plant fitness components (Doak and Morris, 2010), so it may be important to consider multiple plant responses at once (i.e., multiple fitness components or traits).

Biogeographic processes can also shape mating system variation on the landscape. During range expansions, individuals capable of reproduction in the absence of mates or pollinators may be more likely to found new populations (Baker, 1955; Pannell et al., 2015), creating a geographic cline in mating system variation, with a greater degree of self-compatibility or capacity for self-pollination near expanding or recently expanded range edges. Similar patterns might arise in regions where populations turn over frequently, where the ability to reproduce autonomously may be an important trait for individuals that are colonizing empty patches. Gradients in extinction-recolonization dynamics from range centers to edges are hypothesized to create range limits (Holt and Keitt, 2000). Despite these long-standing hypotheses, empirical examinations of mating systems are infrequently carried out at the scale of geographic ranges (with exceptions including Busch 2005; Herlihy and Eckert 2005; Moeller and Geber 2005; Dart et al. 2011; Mimura and Aitken 2007).

In this study, we investigate the relationships among climate, pollinator exclusion, and reproductive fitness components of a winter annual wildflower, *Clarkia pulchella*. In a previous study, we used herbarium specimens to examine relationships between climate, mating system, and reproductive characteristics of this species. We found that summer precipitation was positively correlated with reproductive output and that warm temperatures were correlated with traits indicative of self-pollination (Bontrager and Angert, 2016). Here, we employed field manipulations across the range of *C. pulchella* to examine whether reproductive assurance co-varies with geographic range position and/or climate. *C. pulchella* grows in sites that are very dry during the flowering season, particularly at the northern and southern range edges, so we expected that plants in these regions might have greater capacity to self-pollinate as a means of ensuring reproduction before drought-induced mortality. We therefore predicted that range edge populations would have greater capacity to self-pollinate in the absence of pollinators, and that this geographic pattern would be attributable to climate, in particular, drought stress during the flowering season (summer precipitation and temperature). We anticipated that drought would have opposing effects on reproduction via reproductive assurance vs. fruit production: while drought may prompt increases in reproductive assurance (potentially via plastic responses), it may also limit plant longevity and productivity during the flowering season. Finally, we investigated whether abundance of this species is dependent upon seed production in the previous year.

## Methods

### Study system

*Clarkia pulchella* Pursh (Onagraceae) is a mixed-mating winter annual that grows east of the Cascade Mountains in the interior Pacific Northwest of North America (Fig. 1). The species is found in populations ranging in size from hundreds to thousands of individuals on dry, open slopes in coniferous forest and sagebrush scrub. It is primarily outcrossed by solitary bees (Palladini and Maron, 2013) with a diverse array of other pollinators (MacSwain et al., 1973), but selfing can be facilitated by spatial and temporal proximity of fertile anthers and stigma within flowers.

**Fig. 1.**
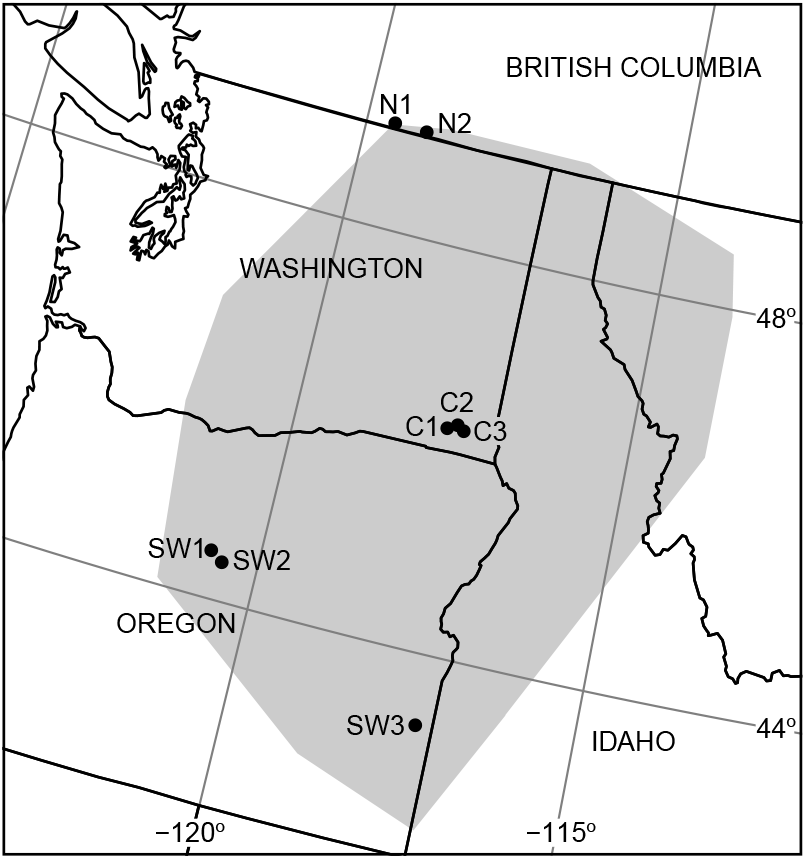
Experimental sites relative to the geographic range of *Clarkia pulchella* (shaded area). N1 and N2 are northern sites; S1, S2, and S3 are southwestern sites, and C1, C2, and C3 are central sites. For geographic coordinates and elevations, see Table S1.

### Plot establishment and monitoring

Experimental plots were established in eight populations on 4-9 June 2015. These sites were located in three regions across the latitudinal range of *Clarkia pulchella*, with two at the species’ northern edge in southern British Columbia (Canada), three in the range center in southeastern Washington (USA), and three in the southwest portion of the species range, in Oregon (USA; Fig. 1, Table S1). Our original intention was to treat the southern and western edges of the range separately and establish three sites at each edge. However, due to difficulty finding populations of sufficient size in sites where we could also obtain permits, we used just two populations in the west and one in the south. Because the climatic similarity among these sites is nearly comparable to that among sites in other regions (Fig. 2, see below for climate data description), we decided to treat them as a single region, the southwest. At each site, 5-8 blocks containing four plots each were marked with 6-inch steel nails, this resulted in a total of 50 blocks and 200 plots in the experiment. Each plot consisted of a 0.8 m^2^ area. Plots were intentionally placed with the goal of obtaining 5-20 individuals per plot, therefore the density in plots was typically higher than the overall site density. If high-density plots have higher-than-average pollinator visitation, our estimates of reproductive assurance could be conservative. Plots were placed closer to other plots in their block than to those in other blocks (with exceptions in two circumstances where low plant density meant very few suitable plot locations were available). Blocks were placed to capture variation in microhabitat characteristics across the site, and their spacing varied depending on the population size and density. Each plot was randomly assigned to one of four factorial treatment groups: control, water addition, pollinator exclusion, or both water addition and pollinator exclusion. Plots receiving water additions were at least 0.5 m away from unwatered plots, except when they were downslope from unwatered plots, in which case they were sometimes closer. Plots receiving pollinator exclusion treatments were tented in bridal-veil mesh with bamboo stakes in each corner and nails tacking the mesh to the ground. Some pollinator exclusion plots had their nets partially removed by wind or cows during the flowering season (n = 13 out of 100 total tented plots), so all analyses were performed without these plots.

**Fig. 2.**
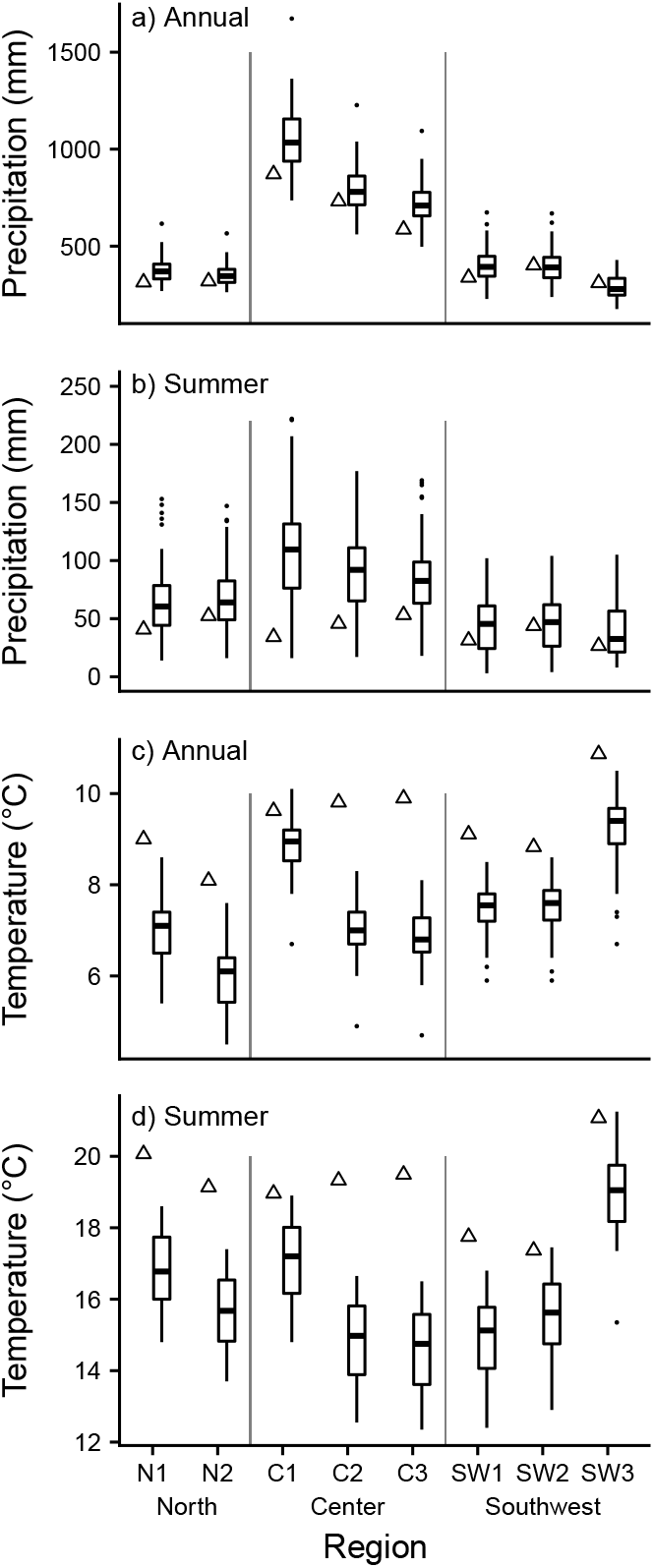
Boxplots summarize a) annual precipitation (mm), b) summer precipitation (mm), c) annual temperature (°C), and d) summer temperature (°C) over a 50-year time window (1963-2012) in each region. Triangles indicate conditions during the experiment. Annual historic values were extracted from ClimateWNA (Wang et al. 2012). Precipitation and temperature during the experiment were downloaded from PRISM (PRISM Climate Group, Oregon State University, prism.oregonstate.edu).

The majority of the summer precipitation in these sites falls in summer storms. In our water addition plots, we added 15 mm of water to each plot (9.6 L per plot) 1-2 times during the flowering season. This approximated the typical precipitation of a summer rainfall event based on data from Wang et al. (2012), and in an average year, would have increased the total summer precipitation in these plots by 30-70%. However, our experiment was conducted during a dry year (Fig. 2), therefore we consider these additions to provide some drought relief, because unwatered plots were already experiencing natural drought. While these treatments measurably increased the soil moisture content immediately following water addition, preliminary analyses showed little effect of water addition on reproduction, and we believe that the timing of amount of water added was not enough to be biologically meaningful. Because of this, we omitted this factor from our final analyses to keep models simple and facilitate interpretation of other factors.

When flowering and fruiting were complete, we counted the number of plants in each plot and the number of fruits on each plant, and estimated the average number of seeds per fruit. The number of plants in each plot ranged from 1-43 (mean = 7.9, median = 7). We counted the number of fruits per plant on every plant in each plot; fruits on *Clarkia pulchella* generally persist on the plant even under pollen limitation (M. Bontrager, personal observation). Plants that had died before producing any flowers were not included in our analyses. Some plants (n = 14, 0.7% of all plants counted) had experienced major damage prior to our final census making fruit counting impossible, so they were assigned the average number of fruits per plant in that plot type at that site for estimation of plot-level seed input, but we excluded them from analyses of fruit counts. Other plants (n = 25, 1.4% of all plants counted) still had flowers at the time of the final census. It was assumed that these flowers would ripen into fruits, so they were included in the fruit counts. When possible, up to four fruits per plot (average number of fruits per plot = 3.67) were collected for seed counting. After counting, seeds were returned to the plots that they were collected from by sprinkling them haphazardly over the plot from a 10 cm height. In 3 of 200 plots, no intact fruits were available for seed counting (all had dehisced), so these plots were excluded from analyses of seed set and plot-level seed input, but included in analyses of fruit counts. To assess the subsequent effects of pollinator limitation on populations in the following year, we revisited plots on 21-24 June and 29-31 July 2016 and counted the number of mature plants present in each. Some plot markers were missing, but we were able to relocate 182 of our 200 plots.

### Climate variable selection

We expect long-term climatic conditions, particularly those that might contribute to drought stress, to influence selection for autonomous selfing. Concurrent work with *C. pulchella* (Bontrager and Angert, 2018) has indicated that fall, winter, and spring growing conditions play a large role in overall plant growth and reproductive output, therefore we considered not only flowering season (June-July) climate variables but also annual temperature and precipitation for inclusion as predictors. We obtained 50-year climate normals (1963-2012) from ClimateWNA (Wang et al., 2012) and climate data during the study from PRISM (PRISM Climate Group, Oregon State University, prism.oregonstate.edu, downloaded 10 October 2016). Our selected set of climatic variables included annual temperature normals (MAT), annual precipitation normals (MAP), summer temperature during the experiment, and summer precipitation during the experiment. Among these, MAT and precipitation during the experiment were correlated (r = −0.84, *P* = 0.0096). A full set of annual and seasonal variable correlations is presented in Table S2.

### Statistical analyses

We used generalized linear mixed effects models (GLMMs) to evaluate the effects of pollinator exclusion, region, and each of the selected climate variables on reproductive assurance and fruits per plant. For each predictor variable of interest (the four climate variables and region), we built a model with a twoway interaction between this variable and pollinator exclusion on both seed counts and fruit counts. We used negative binomial GLMMs for both seeds and fruits, and we included a zero-inflation parameter when modeling seed counts. In all models we included random effects of blocks nested within sites. Because our data do not contain true zero fruit counts (i.e., we did not include plants that did not survive to produce fruits, so all plants in our dataset produced at least one fruit), we subtracted one from all counts of fruits per plant prior to analysis in order to better conform to the assumptions of the negative binomial model. All climate predictors were scaled prior to analyses by subtracting their mean and dividing by their standard deviation. We evaluated the relationship between total plot-level seed production in 2015 and the number of plants present in each plot in summer of 2016 using a GLMM with a negative binomial distribution and random effects of block nested within site. All models were built in R (R Core Team, 2017) using the package glmmTMB (Brooks et al., 2017) and predictions, averaged across random effects, were visualized using the package ggeffects (Lüdecke, 2018).

## Results

### Variation in response to pollinator limitation across the range

In all regions, *Clarkia pulchella* produced fewer seeds in the absence of pollinators (Table 1). In the center and southwest of the range, pollinator exclusion reduced seed set to approximately 30% of control levels (7.2 vs. 24.0 seeds per fruit in the center, 6.4 vs. 19.6 seeds per fruit in the southwest). In the north, pollinator exclusion reduced seed set to about 50% of control levels (14.3 vs. 29.6 seeds per fruit). Climatic or geographic drivers of variation in reproductive assurance were indicated by our models of seeds per fruit when there was a significant interaction between pollinator exclusion and region or pollinator exclusion and a given climate variable. We found that reproductive assurance varied by region, with greater rates of reproductive assurance in northern populations (Fig. 3a, Table 1). Plants in the pollinator exclusion treatment produced about twice as many seeds in the north as in other regions. We did not find any effects of climate on seed production or reproductive assurance (Table 1).

**Fig. 3.**
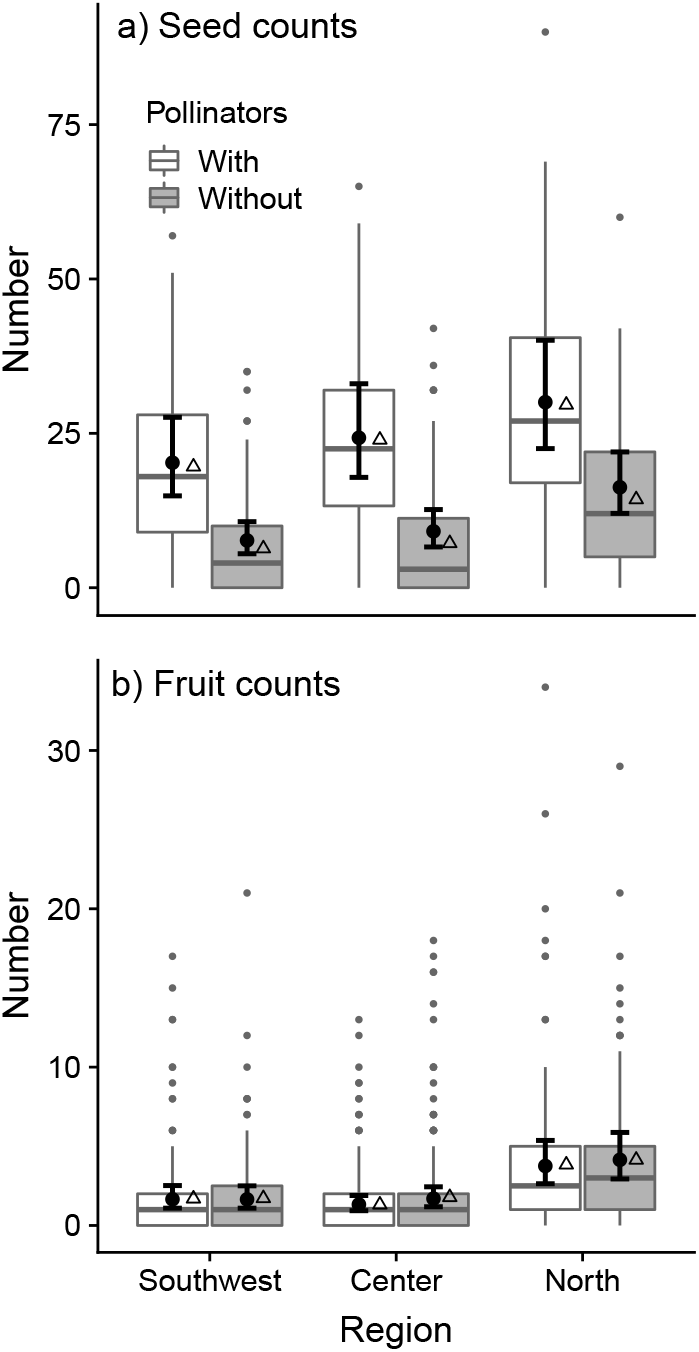
Relationship between a) summer precipitation during the experiment (2015) and b) mean annual temperature (1963-2012) on per-plant fruit production with and without pollinators. Lines represent model predictions; shaded areas represent 95% confidence intervals. Each pair of stacked dots represent raw means of plants with (black) and without pollinators (grey) in a site.

**Table 1.** Effects of pollinator exclusion, region, and climate on seed set per fruit. Estimates, standard errors, and *P*-values are from zero-inflated negative binomial GLMMs. Effects of being in the northern or southwestern region are expressed relative to central populations. Significant main effects and interactions are indicated with bold font.

| Climate/region predictor |  | Climate/region |  |  | Pollinator exclusion |  |  | Climate/region x pollinator exclusion |  |  |
| --- | --- | --- | --- | --- | --- | --- | --- | --- | --- | --- |
| | | $\beta$ | SE | $P$-value | $\beta$ | SE | $P$-value | $\beta$ | SE | $P$-value |
| Region | North | 0.219 | 0.155 | 0.157 | **-0.987** | **0.112** | **< 0.001** | **0.371** | **0.159** | **0.020** |
|  | Southwest | -0.217 | 0.139 | 0.119 |  |  |  | 0.098 | 0.153 | 0.523 |
| Mean annual precipitation |  | 0.046 | 0.098 | 0.635 | **-0.834** | **0.066** | **< 0.001** | -0.111 | 0.064 | 0.086 |
| Mean annual temperature |  | -0.084 | 0.090 | 0.348 | **-0.827** | **0.066** | **< 0.001** | -0.038 | 0.062 | 0.541 |
| Summer precipitation (2015) |  | 0.031 | 0.096 | 0.747 | **-0.825** | **0.066** | **< 0.001** | -0.019 | 0.063 | 0.763 |
| Summer temperature (2015) |  | 0.037 | 0.093 | 0.688 | **-0.840** | **0.067** | **< 0.001** | 0.105 | 0.065 | 0.105 |

### Response of patch density to seed production in the previous year

Across sites, there was a positive relationship between the number of seeds produced in a plot in 2015 and the number of adult plants present in 2016 (*P* < 0.0001, *β* = 0.00044, SE = 0.000061). This is not simply a result of plots with large numbers of plants in 2015 being similarly dense in 2016, because seed input was decoupled from plant density in 2015 by the pollinator exclusion treatments. The effect of seed input remained significant (*P* < 0.0001) when the number of plants in 2015 was included in the model as a covariate (results not shown).

### Variation in fruit production across the range

Plants in the north produced more fruits (on average 4.0, compared to 1.5 and 1.7 in the center and southwest, respectively; Table 2, Fig. 3b). This regional trend could be due to the relatively lower normal annual temperatures in the northern sites (Fig. 2), the effects of which are discussed below. Pollinator exclusion tended to result in a slight increase in fruit production, possibly due to reallocation of resources within a plant in order to produce more flowers when ovules are left unfertilized (Table 2). This effect was small—plants in plots without pollinators produced an additional 0.4 fruits, on average (Fig. 3b).

**Table 2.** Effects of pollinator exclusion, region, and climate on fruit number. Estimates, standard errors, and *P*-values are from negative binomial GLMMs. Effects of being in the northern or southwestern region are expressed relative to central populations. Significant main effects and interactions are indicated with bold font.

| Climate/region predictor |  | Climate/region |  |  | Pollinator exclusion |  |  | Climate/region x pollinator exclusion |  |  |
| --- | --- | --- | --- | --- | --- | --- | --- | --- | --- | --- |
| | | $\beta$ | SE | $P$-value | $\beta$ | SE | $P$-value | $\beta$ | SE | $P$-value |
| Region | North | **1.156** | **0.442** | **0.009** | **0.302** | **0.096** | **0.002** | -0.272 | 0.146 | 0.063 |
|  | Southwest | 0.001 | 0.399 | 0.997 |  |  |  | -0.153 | 0.151 | 0.309 |
| Mean annual precipitation |  | -0.126 | 0.236 | 0.594 | **0.178** | **0.062** | **0.004** | 0.020 | 0.064 | 0.752 |
| Mean annual temperature |  | **-0.454** | **0.148** | **0.002** | **0.154** | **0.063** | **0.014** | -0.118 | 0.066 | 0.072 |
| Summer precipitation (2015) |  | 0.281 | 0.206 | 0.172 | **0.148** | **0.062** | **0.017** | **0.211** | **0.068** | **0.002** |
| Summer temperature (2015) |  | -0.253 | 0.201 | 0.209 | **0.174** | **0.062** | **0.005** | -0.033 | 0.066 | 0.617 |

We found that the effects of pollinator exclusion on fruit production depended upon the amount of summer precipitation during the experiment (Table 2, Fig. 4). Fruit production was higher in wetter sites, and pollinator-excluded plants that were in the wettest sites showed a greater positive effect of pollinator exclusion on fruit production (Table 2). However, it should be noted that while both the main effect of climate and its interaction with pollinator exclusion were significant, the difference between plots with and without pollinators in wetter sites did not appear to be particularly strong, and when visualized the confidence intervals were largely overlapping (Fig. 4A). We also found a main effect of mean annual temperature (MAT) on fruit production (Table 2). Fruit production was higher in cooler sites (Fig. 4B). Disentangling these two climatic drivers of increased fruit production is not possible with this dataset, however, because summer precipitation during the experiment was negatively correlated with normal MAT. Therefore, it could have been either higher water resources during flowering or cooler temperatures over the growing season that resulted in increased fruit production. It is worth noting, however, that summer temperature during the experiment was not correlated with either of these variables, so if temperature was the driver of this pattern, it was likely because of temperature effects on earlier life-history stages.

**Fig. 4.**
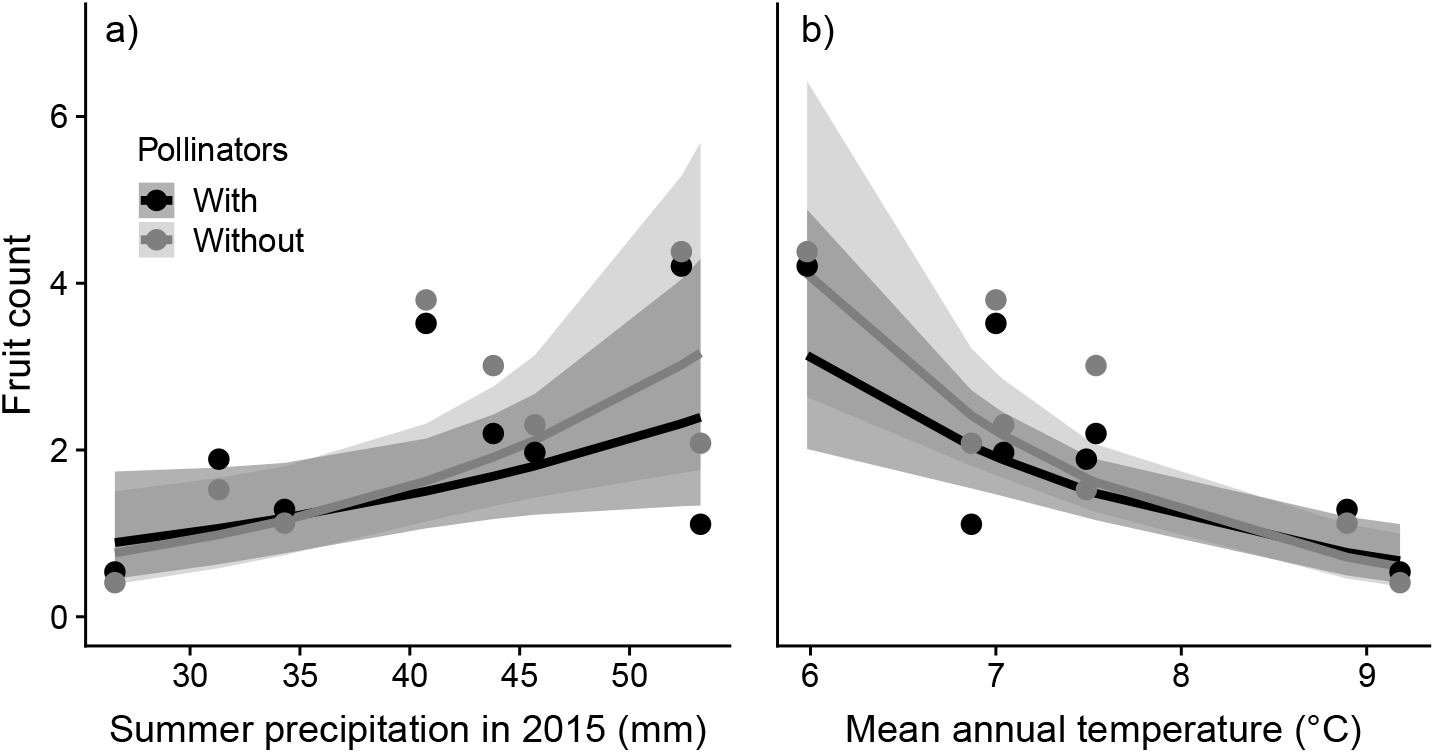
Relationship between a) summer precipitation during the experiment (2015) and b) mean annual temperature (1963-2012) on per-plant fruit production with and without pollinators. Lines represent model predictions; shaded areas represent 95% confidence intervals. Each pair of stacked dots represent raw means of plants with (black) and without pollinators (grey) in a site.

## Discussion

Pollinator exclusion in eight populations of *Clarkia pulchella* revealed increased autonomous reproductive assurance in populations in the northern part of the species’ range, as compared to the center or southwest. Plants in the northern part of the species’ range also produced more fruits. Fruit production was higher in sites that are cooler or that received higher amounts of precipitation during the experiment. Plants also produced slightly more fruits in response to pollinator exclusion, however, this reallocation was not, in general, large enough to offset the reduction in seed production caused by pollen limitation.

### Reproductive assurance is driven by geography rather than climate

Pollinator limitation reduced reproduction across the range of *C. pulchella*. Contrary to our prediction, we did not observe decreased reproductive assurance in sites with high summer precipitation during the experiment. Increased reproductive assurance was only apparent at the northern range edge (Fig. 3a) and was not associated with any of the climate variables that we considered. In light of this, we suggest that for this species, reproductive assurance is better explained by the latitudinal position of populations than by any single climate variable. The locations of our northern populations were covered by the Cordilleran ice sheet during the last glacial maximum; the patterns we see could be the result of a post-glacial range expansion, in which the founders of these northern populations were individuals who had a greater capacity for autonomous reproduction (Baker, 1955). It is possible that during colonization there is a low probability of pollinators foraging on a novel plant species and moving conspecific pollen between sparse individuals. Reproductive assurance has evolved in other species when populations have experienced historic bottlenecks (Busch, 2005), and contrasts of species’ range sizes indicate that species capable of autonomous self-pollination have a greater ability to colonize new sites (Randle et al., 2009). While latitude is not a strong predictor of among-species variation in mating system (Moeller et al., 2017), within-species variation may be more closely tied to postglacial colonization routes.

An alternative possibility is that our northern sites are distinct because they differ in community composition from sites in other parts of the range. Our experimental sites may differ in their regional suite of pollinators. Geographic variation in pollinator communities can be related to floral traits (Nattero and Cocucci, 2007) and plant reproductive success (Gómez et al., 2007) and has been linked to mating system varition in closely related taxa (Moeller, 2006). Alternatively, or additionally, our sites could differ in the co-occurring plant community. In a related species, co-occurrence with congeners relaxes selection for self-pollination, presumably by alleviating Allee effects (Moeller and Geber, 2005). Contrastingly, plant species that compete for pollinators can reduce deposition of conspecific pollen on *C. pulchella* (Palladini and Maron, 2013). It is possible that populations in the northern portion of the range have adapted to a different pollination environment caused by overlap with different plant species.

Across the range, adult plant density was positively correlated with seed production in the previous year. Because our pollinator exclusion treatment led to plot-level seed input being decoupled from the number of plants in 2015 (data not shown), we can attribute differences in 2016 plant density to seed input, rather than to patch quality. Seed production is important enough to have an effect on subsequent density despite differences between plots in the availability of germination sites or the probability of survival to flowering. This, in combination with the consistent negative reproductive response to pollinator exclusion, indicates that populations would likely be negatively impacted by disruption of pollinator service.

### Reallocation to flower and fruit production under pollen limitation

Either cool temperatures during the growing season, high summer precipitation, or a combination of the two increase overall fruit production. Germination of *C. pulchella* is inhibited under warm temperatures (Lewis, 1955), so plants in sites with cooler fall temperatures could have earlier germination timing and develop larger root systems, giving them access to more resources during the flowering season. *Clarkia pulchella* individuals appear to be capable of reallocating some resources to flower production when pollen is limited (Table 2). Our finding of a modest amount of reallocation under pollinator exclusion contrasts with work in another *Clarkia* species, *C. xantiana* ssp. *parviflora*, which found that individuals do not reallocate resources based on the quantity of pollen received (Briscoe Runquist and Moeller, 2013). These contrasting results can potentially be explained by two factors. First, the focal species of our study produces buds continuously over the flowering season, while *C. xantiana* ssp. *parviflora* produces nearly all of its buds at the beginning of the flowering season, leaving individuals little opportunity to respond to the pollination environment (Briscoe Runquist and Moeller, 2013). Second, their study investigated differences between plants under natural pollination conditions and plants receiving supplemental pollen, while we compared plants under natural pollination and plants under strong pollen limitation. These differences in direction and magnitude of the treatments imposed may affect the degree to which a plant reallocation response can be detected. An alternative explanation for the apparent resource reallocation is that our pollinator exclusion tents protected plants from herbivores that might have removed fruits in the control plots. While herbivory of individual fruits (rather than entire plants) appears rare (M. Bontrager, personal observation), we can not rule out the possibility of an herbivore effect.

### Conclusions and future directions

Populations of *Clarkia pulchella* from across the species’ range are reliant on pollinator service to maintain high levels of seed production, which our data show to be an important demographic component that influences abundance for this species. Our data support the hypothesis that populations in areas of the range that have undergone post-glacial expansion may have elevated levels of reproductive assurance, but alternative drivers of this pattern remain plausible. Future work should explore these drivers, and could begin by examining geographic variation in the phenology, abundance, and composition of pollinator communities. In order to better understand how *C. pulchella* might respond to changes in pollinator service, future work should measure the capacity of populations to evolve higher rates of self-pollination in the absence of pollinators.

## Acknowledgements

We would like to thank B. Harrower, R. Germain, and members of the Angert lab for their thoughtful comments on this project. We also greatly appreciate helpful comments from two anonymous reviewers. E. Fitz assisted with fieldwork. Permission to work in our field sites was granted by British Columbia Parks, Umatilla National Forest, Ochoco National Forest, and the Vale District Bureau of Land Management. MB was supported by a University of British Columbia Four-year Fellowship, and this work was also supported by a Natural Sciences and Engineering Research Council of Canada Discovery Grant to ALA.

## Data accessibility

Data and code are available on Github at https://github.com/meganbontrager/clarkia-reproductive-assurance. Data and code are archived on Zenodo (doi: 10.5281/zenodo.2597693).

---

**Table S1.** Geographic data for experimental sites. Coordinates are given in decimal degrees. Abbreviations match Fig. 1.

| Name | Abbreviation | Latitude | Longitude | Elevation (m) |
| --- | --- | --- | --- | --- |
| Southwest 1 | SW1 | 44.47 | -120.71 | 1128 |
| Southwest 2 | SW2 | 44.38 | -120.52 | 1134 |
| Southwest 3 | SW3 | 43.30 | -117.27 | 1043 |
| Center 1 | C1 | 46.24 | -117.74 | 1022 |
| Center 2 | C2 | 46.28 | -117.60 | 1457 |
| Center 3 | C3 | 46.24 | -117.49 | 1445 |
| North 1 | N1 | 49.05 | -119.56 | 842 |
| North 2 | N2 | 49.04 | -119.05 | 866 |

**Table S2.** Pearson correlation coefficients among precipitation and temperature variables associated with experimental sites. For **(A)** precipitation, **(B)** temperature, and **(C)** precipitation and temperature, correlations are shown between annual, fall (September-November), winter (December-February), spring (March-May), and summer (June-July, because all plants senesce before August). Normal values were calculated over 50 years (1963-2012), while 2014-2015 values are from the growing season of plants in the experiment. Normal climate data is from ClimateWNA (Wang et al., 2012) and 2014-2015 variables are from PRISM (PRISM Climate Group, Oregon State University, prism.oregonstate.edu). Variables used in models and their correlations are indicated in bold text.

| A. Temperature |  |  |  |  |  |  |  |  |  |
| --- | --- | --- | --- | --- | --- | --- | --- | --- | --- |
|  | **MAT**<br>**(normal)** | Fall<br>temp.<br>(normal) | Winter<br>temp.<br>(normal) | Spring<br>temp.<br>(normal) | Summer<br>temp.<br>(normal) | MAT<br>(2014-15) | Fall<br>temp.<br>(2014) | Winter<br>temp.<br>(2014-15) | Spring<br>temp.<br>(2015) |
| Fall temp. (normal) | 0.97 |  |  |  |  |  |  |  |  |
| Winter temp. (normal) | 0.69 | 0.83 |  |  |  |  |  |  |  |
| Spring temp. (normal) | 0.86 | 0.73 | 0.23 |  |  |  |  |  |  |
| Summer temp. (normal) | 0.72 | 0.54 | 0.01 | 0.94 |  |  |  |  |  |
| MAT (2014-2015) | 0.71 | 0.68 | 0.62 | 0.44 | 0.48 |  |  |  |  |
| Fall temp. (2014) | 0.70 | 0.75 | 0.82 | 0.30 | 0.26 | 0.95 |  |  |  |
| Winter temp. (2014-2015) | 0.60 | 0.74 | 0.95 | 0.11 | -0.04 | 0.71 | 0.90 |  |  |
| Spring temp. (2015) | 0.28 | 0.05 | -0.38 | 0.57 | 0.78 | 0.40 | 0.08 | -0.35 |  |
| **Summer temp. (2015)** | **0.25** | 0.05 | -0.24 | 0.40 | 0.65 | 0.59 | 0.30 | -0.14 | 0.93 |
| B. Precipitation |  |  |  |  |  |  |  |  |  |
|  | **MAP**<br>**(normal)** | Fall<br>precip.<br>(normal) | Winter<br>precip.<br>(normal) | Spring<br>precip.<br>(normal) | Summer<br>precip.<br>(normal) | MAP<br>(2014-15) | Fall<br>precip.<br>(2014) | Winter<br>precip.<br>(2014-15) | Spring<br>precip.<br>(2015) |
| Fall precip. (normal) | 1.00 |  |  |  |  |  |  |  |  |
| Winter precip. (normal) | 1.00 | 1.00 |  |  |  |  |  |  |  |
| Spring precip. (normal) | 0.99 | 0.99 | 0.98 |  |  |  |  |  |  |
| Summer precip. (normal) | 0.92 | 0.88 | 0.88 | 0.89 |  |  |  |  |  |
| MAP (2014-2015) | 0.99 | 0.99 | 0.99 | 0.98 | 0.88 |  |  |  |  |
| Fall precip. (2014) | 0.98 | 0.98 | 0.98 | 0.97 | 0.87 | 0.98 |  |  |  |
| Winter precip. (2014-2015) | 0.98 | 0.98 | 0.98 | 0.98 | 0.86 | 1.00 | 0.98 |  |  |
| Spring precip. (2015) | 0.96 | 0.97 | 0.96 | 0.98 | 0.82 | 0.98 | 0.95 | 0.99 |  |
| **Summer precip. (2015)** | **0.13** | 0.10 | 0.11 | 0.05 | 0.38 | 0.11 | 0.04 | 0.09 | 0.01 |

C. Precipitation and temperature
|  | **MAT**<br>**(normal)** | Fall<br>temp.<br>(normal) | Winter<br>temp.<br>(normal) | Spring<br>temp.<br>(normal) | Summer<br>temp.<br>(normal) | MAT<br>(2014-15) | Fall<br>temp.<br>(2014) | Winter<br>temp.<br>(2014-15) | Spring<br>temp.<br>(2015) | **Summer**<br>**temp.**<br>**(2015)** |
| --- | --- | --- | --- | --- | --- | --- | --- | --- | --- | --- |
| **MAP (normal)** | **0.20** | 0.30 | 0.47 | -0.05 | -0.19 | 0.23 | 0.31 | 0.37 | -0.18 | **-0.08** |
| Fall precip. (normal) | 0.23 | 0.34 | 0.53 | -0.05 | -0.21 | 0.22 | 0.33 | 0.41 | -0.24 | -0.14 |
| Winter precip. (normal) | 0.22 | 0.33 | 0.53 | -0.06 | -0.22 | 0.22 | 0.33 | 0.42 | -0.25 | -0.15 |
| Spring precip. (normal) | 0.30 | 0.37 | 0.52 | 0.03 | -0.10 | 0.34 | 0.41 | 0.42 | -0.09 | 0.03 |
| Summer precip. (normal) | -0.08 | -0.04 | 0.09 | -0.17 | -0.21 | 0.04 | 0.03 | 0.01 | 0.01 | 0.09 |
| MAP (2014-2015) | 0.25 | 0.34 | 0.51 | -0.02 | -0.16 | 0.28 | 0.37 | 0.43 | -0.19 | -0.07 |
| Fall precip. (2014) | 0.28 | 0.37 | 0.50 | 0.04 | -0.13 | 0.21 | 0.30 | 0.38 | -0.18 | -0.12 |
| Winter precip. (2014-2015) | 0.28 | 0.37 | 0.53 | 0.00 | -0.14 | 0.31 | 0.40 | 0.46 | -0.18 | -0.06 |
| Spring precip. (2015) | 0.34 | 0.43 | 0.61 | 0.02 | -0.12 | 0.40 | 0.49 | 0.53 | -0.17 | -0.02 |
| **Summer precip. (2015)** | **-0.84** | -0.80 | -0.51 | -0.79 | -0.67 | -0.50 | -0.48 | -0.39 | -0.28 | **-0.17** |

